# Single-cell RNA-seq of the stromal vascular fraction of adipose tissue reveals lineage-specific changes in cancer-related lymphedema

**DOI:** 10.1101/2020.09.27.315911

**Authors:** Xuanyu Liu, Meng Yuan, Qinqin Xiang, Wen Chen, Zhujun Li, Jie Chen, Jiuzuo Huang, Nanze Yu, Xiao Long, Zhou Zhou

## Abstract

Lymphedema is a chronic tissue edema that frequently occurs following lymph node resection for cancer treatment, and is characterized by progressive swelling, chronic inflammation, excessive fibrosis and adipose deposition in the affected limbs. We still lack targeted medical therapies for this disease due to the incomplete understanding of the mechanism underlying the pathogenesis. Here, we performed single-cell RNA-seq of 70,209 cells of the stromal vascular fraction (SVF) of subcutaneous adipose tissue from patients with cancer-related lymphedema and healthy donors. Unbiased clustering revealed 21 cell clusters, which were assigned to 10 cell lineages. One of the four ASC subpopulations, c3, was significantly expanded in lymphedema, which may be related to the fibrosis and pathologic mineralization of adipose tissues in lymphedema. Dysregulated pathways and genes of ASCs in lymphedema were identified through gene set enrichment analysis and differential regulatory network analysis, which reflect the pathophysiological changes in ASCs in lymphedema: enhanced fibrosis, mineralization and proliferation as well as compromised immunosuppression capacity. In addition, we characterized the three subpopulations of macrophages, and found that the adipose tissue of lymphedema displayed immunological dysfunction characterized by a striking depletion of anti-inflammatory macrophages, i.e., *LYVE^+^* resident-like macrophages. Cell-cell communication analysis revealed a perivascular ligand-receptor interaction module among ASCs, macrophages and vascular endothelial cells in adipose tissue. Communication changes for ASCs in lymphedema were identified. For example, PDGFD-PDGFR complex interactions were significantly enhanced between a number of lineages and ASCs, reflecting the role of PDGFD signaling in the pathophysiological changes in ASCs. Finally, we mapped the previously reported candidate genes predisposing to cancer-related lymphedema to cell subpopulations in the SVF, and found that *GJC2*, the most likely causal gene was highly expressed in the lymphedema-associated ASC subpopulation c3. In summary, we provided the first comprehensive analysis of cellular heterogeneity, lineage-specific regulatory changes and intercellular communication alterations of the SVF in adipose tissues from cancer-related lymphedema at a single-cell resolution. The lymphedema-associated cell subpopulations and dysregulated pathways may serve as potential targets for medical therapies. Our large-scale dataset constitutes a valuable resource for further investigations of the mechanism of cancer-related lymphedema.

## Introduction

Lymphedema is a chronic tissue edema that results from lymphatic drainage disorders due to intrinsic fault (primary lymphedema) or damage (secondary lymphedema) to the lymphatic system (Lawenda et al., 2009). Secondary lymphedema is the most prevalent form and frequently occurs following lymph node resection for cancer treatment, i.e., cancer-related lymphedema (Shaitelman et al., 2015). Up to 20% of women develop this condition following treatment for breast cancer (DiSipio et al., 2013). Lymphedema is characterized by progressive swelling, chronic inflammation, excessive fibrosis and adipose deposition in the affected limbs (Zampell et al., 2012a). Lymphedema usually exerts a significant physical and psychological burden on cancer survivors and severely affects their quality of life; however, the clinical treatment remains palliative (Shaitelman et al., 2015). We still lack effective therapies, in particular, targeted medical therapies, for the treatment or prevention of this complication, which is partially due to the incomplete understanding of the cellular mechanism of pathogenesis.

Adipose tissue is not simply a container of fat, but an endocrine organ, which is composed of multiple types of cells, such as adipose-derived stromal/stem/progenitor cells (ASCs), adipocytes, vascular cells (e.g., vascular endothelial cells and pericytes) and immune cells (e.g., macrophages and lymphocytes) (Vijay et al., 2020). All nonadipocyte cells are known as the stromal vascular fraction (SVF), which can be isolated through enzymatic digestion (Ramakrishnan and Boyd, 2018). Lymphatic fluid stasis in the limbs of patients with lymphedema will ultimately result in increased subcutaneous adipose tissue volume and excess adipose deposition, which may lead to further deterioration of the lymphatic system (Mehrara and Greene, 2014). Previous studies have found significant alterations in the SVF of subcutaneous adipose tissue in lymphedema with regard to cellular composition, proliferation and differentiation capacity, which reflects the role of SVF changes in the pathophysiology of lymphedema (Aschen et al., 2012; Januszyk et al., 2013; Tashiro et al., 2017; Zampell et al., 2012b). However, previous studies generally rely on the expression of a limited number of marker genes and have focused on a few cell lineages. We still lack a comprehensive and accurate understanding of the alterations of adipose tissue in lymphedema.

Recent technical advances in single-cell RNA-seq have enabled the transcriptomes of tens of thousands of cells to be assayed at single-cell resolution (Zheng et al., 2017). Compared with the averaged expression of genes from a mixed cell population obtained by bulk RNA-seq, large-scale single-cell RNA-seq allows unbiased cellular heterogeneity dissection and regulatory network construction at an unprecedented scale and resolution (Kulkarni et al., 2019). Single-cell RNA-seq is therefore emerging as a powerful tool for understanding the cellular and molecular mechanisms of pathogenesis in a variety of diseases such as pulmonary fibrosis (Reyfman et al., 2019) and lupus nephritis (Der et al., 2019). Single-cell RNA-seq has also been applied to dissect the heterogeneity of the SVF in mice (Burl et al., 2018; Schwalie et al., 2018) and humans (Vijay et al., 2020). However, to our knowledge, few studies have been performed to explore the alterations in the SVF under a diseased condition, for example, lymphedema, at a single-cell resolution.

In this study, we performed single-cell RNA-seq of 70,209 cells of the SVF of subcutaneous adipose tissue from patients with cancer-related lymphedema and healthy donors. We aimed to identify cell lineages or subpopulations associated with lymphedema, lineage-specific regulatory changes and intercellular communication alterations in adipose tissue from lymphedema.

## Results

### Single-cell RNA-seq reveals cellular diversity and heterogeneity of the SVF of subcutaneous adipose tissue in patients with cancer-related lymphedema

To unbiasedly dissect the cellular heterogeneity of the SVF of adipose tissue in healthy and diseased conditions (cancer-related lymphedema), we obtained subcutaneous adipose tissue specimens from the affected thighs of five patients with severe lymphoedema (stage III; the CASE group) following surgical intervention for cervical cancer. As a control group, liposuction specimens from the thighs of four healthy female donors were also collected (Figure 1A; Table S1). After SVF isolation, all the samples were subjected to single-cell transcriptomic sequencing. Following stringent quality filtering, we ultimately obtained transcriptomes of 70,209 cells (CASE: 41,274 cells; CTRL: 28,935 cells). Unbiased clustering revealed 21 clusters (Figure 1B). Based on hierarchical clustering (Figure 1C) and established lineage-specific marker genes (Figure 1D), we assigned these clusters to 10 cell lineages. The representative molecular signatures of these clusters are shown in Figure 1E and Table S2.

**Figure 1.**
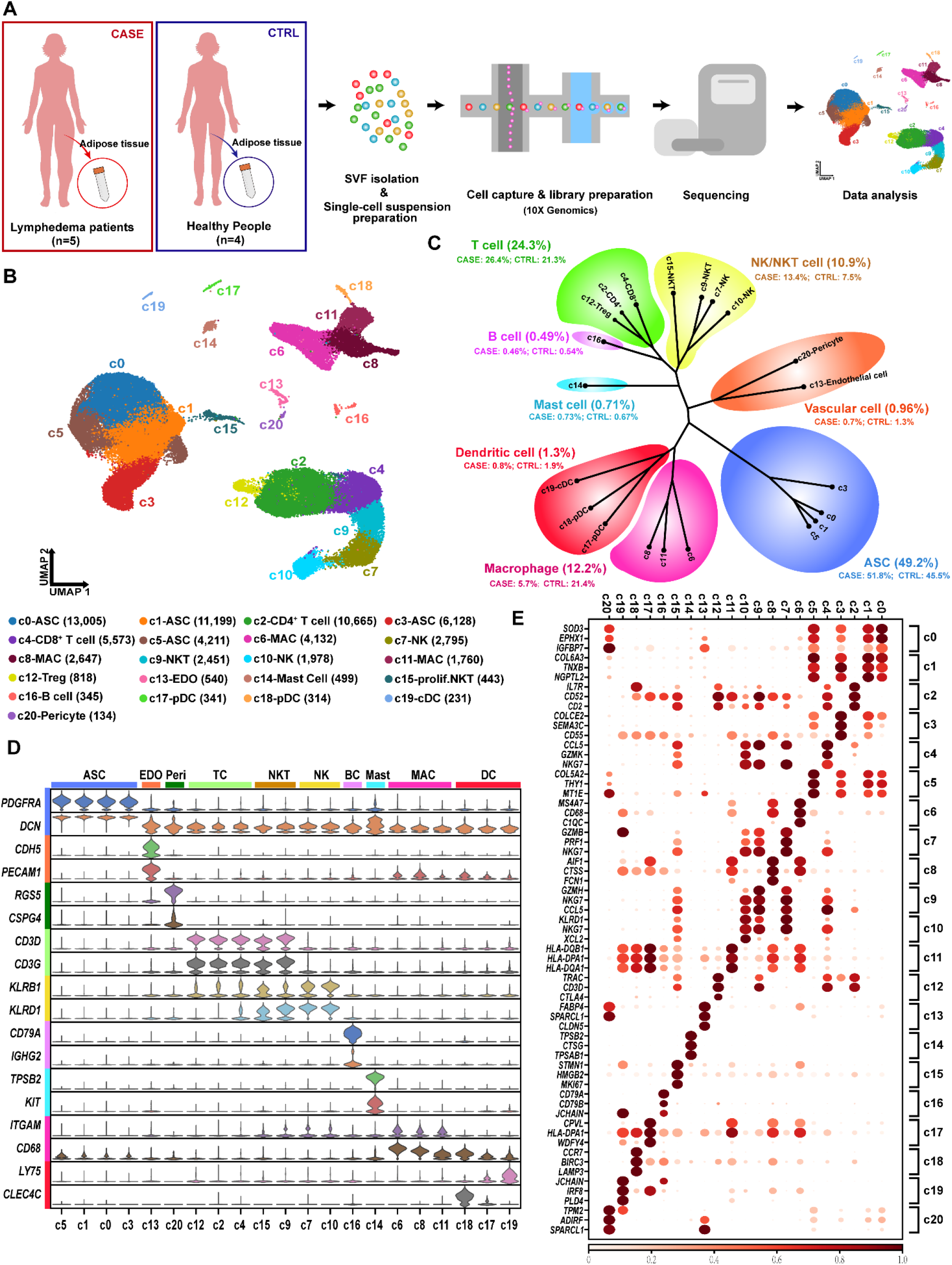
Single-cell RNA-seq reveals cellular diversity and heterogeneity of the SVF of adipose tissue in patients with cancer-related lymphedema. **(A)** Schematic representation of the experimental procedure. Five patients with cancer-related lymphedema (the CASE group) and four healthy people were recruited in this study. Liposuction specimens from the thighs were collected during surgery. **(B)** Unbiased clustering of 70,209 cells revealed 21 cellular clusters. Clusters are distinguished by different colors. The number in parentheses represents the cell count. **(C)** Hierarchical clustering of the clusters based on the average expression of the 2,000 most variable genes. **(D)** Expression of the established marker genes for each lineage in each cluster. **(E)** Representative molecular signatures for each cell cluster. The area of the circles indicates the proportion of cells expressing the gene, and the color intensity reflects the expression intensity. ASC: adipose-derived stromal/stem/progenitor cell; cDC: conventional dendritic cell; EDO: endothelial cell; MAC: macrophage; NK: natural killer cell; NKT: natural killer T cell; prolif.NKT: proliferative nature killer T cell; pDC: plasmacytoid dendritic cell.

The ASC lineage (marked by *PDGFRA* and *DCN*) (Guerrero-Juarez et al., 2019), including c0, c1, c3 and c5, accounted for a large proportion (49.2%) of the SVF (Figure 1C), which is comparable with that (55%) reported previously (Vijay et al., 2020). A large and diverse population of immune cells (49.9%) were found, including both myeloid cells and lymphocytes. The dominant lineage of myeloid cells was macrophages (marked by *ITGAM* and *CD68*) (Singhal et al., 2019), which included three subpopulations, i.e., c6, c8 and c11. Two other types of myeloid cells were mast cells (marked by *TPSB2* and *KIT*) (Vieira Braga et al., 2019) and dendritic cells (DCs). The DCs encompassed clusters of conventional dendritic cells (cDCs; c19; marked by *LY75*) and plasmacytoid dendritic cells (pDCs; c18 and c17; marked by *CLEC4C*) (Merad et al., 2013). The lymphocytes detected included T cells (c2, c4, and c12; marked by *CD3D* and *CD3G*) (Guo et al., 2018), B cells (c16; marked by *CD79A* and *IGHG2*) (Hu et al., 2017), natural killer (NK) cells (c7 and c10; marked by *KLRB1* and *KLRD1*) (Xu et al., 2011) and natural killer T (NKT) cells (c9 and c15; expressing both NK and T cell markers). Detailed analysis revealed that both c2 and c12 belonged to CD4^+^ helper T cells (marked by *CD4* and *IL7R;* Figure S1). Cluster c12 also exhibited expression of *CTLR4* and *FOXP3* (Figure S1), thus representing a cluster of regulatory T cells (Treg cells) (Li et al., 2015). Cluster c4 was a cluster of CD8^+^ T cells, reflected by high expression of *CD8A* and *CD8B* (Figure S1). The NKT cluster c15 expressed high levels of proliferation markers such as *MKI67* and *TOP2A*, thus representing proliferative NKT cells, whereas the NKT cluster c9 belonged to nonproliferative NKT cells (Figure S1). In addition, we identified vascular cells including endothelial cells (c13; marked by *CDH5* and *PECAM1*) (Kalucka et al., 2020) and pericytes (c20; marked by *RGS5* and *CSPG4*) (Holm et al., 2018). Together, single-cell analysis reveals previously unrecognized cellular diversity and heterogeneity of the SVF of subcutaneous adipose tissue in lymphedema.

### Differential proportional analysis reveals significantly expanded or contracted cell lineages associated with cancer-related lymphedema

Cell lineages that greatly change in relative proportion are probably associated with the pathogenesis of the disease. Visualization of the cellular density revealed dramatic changes in the relative proportions of multiple lineages, including ASCs, macrophages and lymphocytes (Figure 2A). To determine whether the proportional change was expected by chance, we performed a permutation-based statistical test (differential proportion analysis; DPA) as described previously (Farbehi et al., 2019). As shown in Figure 2B, the ASCs were significantly expanded (Bonferroni-corrected p-value < 0.01), which suggests enhanced proliferation or differentiation of ASCs in lymphedema. Indeed, we observed significantly higher cycling scores for ASCs in CASE versus CTRL (Wilcoxon rank sum test p-value = 4.916E-09; Figure S2). Strikingly, lymphocyte lineages (T cells, NK and NKT cells) were significantly expanded, whereas the myeloid lineages (macrophages and DCs) were significantly contracted (Bonferroni-corrected p-value < 0.05; Figure 2B). This result may reflect enhanced adaptive immunity and exhausted innate immunity at this severe stage of lymphedema. Further analysis at the cluster level revealed significantly expanded subpopulations, including c2 CD4^+^ T cells; c3 ASCs, c7 NK cells and c9 NKT cells, reflecting a strong association of these subpopulations with pathogenesis (Figure 2C and 2D). The three macrophage subpopulations, especially cluster c6, were greatly contracted. Given the results above and the relatively large cellular proportion, our study focused on the ASC and macrophage lineages, which may play dominant role in the pathogenesis and could potentially serve as cellular targets for medical intervention.

**Figure 2.**
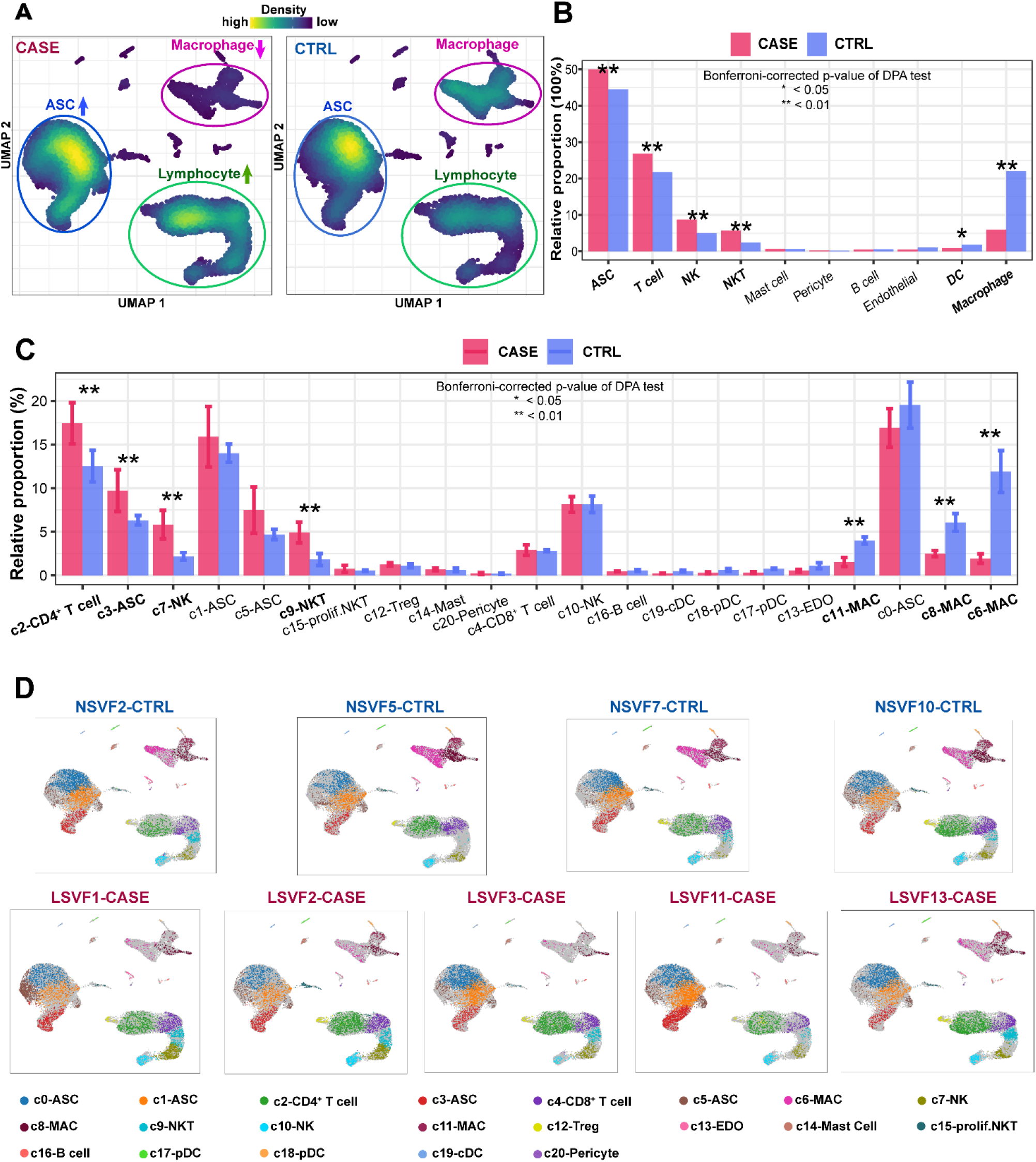
Differential proportional analysis reveals significantly expanded or contracted cell lineages associated with cancer-related lymphedema. **(A)** Visualization of the cellular density reveals dramatic changes in the proportions of multiple cell lineages in CASE versus CTRL. Cells were randomly sampled for equal numbers in the CASE (n= 28,935) and CTRL (n= 28,935) groups in this analysis. **(B)** Significantly expanded or contracted cell lineages. **(C)** Significantly expanded or contracted cell clusters. **(D)** The distribution of cells for each cluster in each individual. In B and C, a permutation-based statistical test (differential proportion analysis; DPA) was performed. A Bonferroni-corrected p-value < 0.05 was considered to be statistically significant.

### Heterogeneity of ASCs in the SVF of adipose tissue unraveled by single-cell analysis

We examined the expression of marker genes normally used for identifying freshly isolated or cultured ASCs (Figure 3A). Consistent with our knowledge (Suga et al., 2009), *CD34*, a marker for freshly isolated ASCs in the SVF, is highly expressed in all ASC subpopulations. The ASCs expressed positive markers for the definition of cultured ASCs (e.g., *CD105, CD73, CD90, CD59, CD44* and CD29) and generally lacked expression of negative markers (e.g., *CD45, CD14, CD11b, CD19* and *CD79A*) (Dominici et al., 2006; Gimble et al., 2007). Notably, we found that some ASCs, particularly in cluster c5, expressed MHC class II genes (e.g., *HLA-DRA, HLA-DRB1* and *HLA-DRB5*), suggesting that these cells had antigen-presenting functions. This finding agrees with the notion that antigen-presenting functions could be induced in inflammatory or diseased states for ASCs, albeit the fact that they are not natural antigen-presenting cells (Liu et al., 2017). Next, we found that the four subpopulations had distinct expression profiles (Figure 3B; Table S3). Cluster c0 expressed high levels of adipose stem cell or preadipocyte markers such as *CXCL14, APOD, APOE, MGP* and *WISP2* (Vijay et al., 2020). The gene signature of c0 was enriched with the Gene Ontology (GO) term “positive regulation of hemostasis” (representative genes: *CD36, F3* and *SELENOP;* Figure 3C). In line with these results, subpopulation-specific regulon analysis using SCENIC (Aibar et al., 2017) identified *PPARG* and *CEBPA*, the known master TFs in adipogenesis (Cristancho and Lazar, 2011), as c0-specific key regulators (Figure 3D). Notably, c3, a lymphedema-associated ASC subpopulation based on the DPA above (Figure 2C), showed high expression of genes specifically expressed by chondrocytes (e.g., *PRG4*) (Kozhemyakina et al., 2015), and its molecular signature was enriched with GO terms such as “collagen fibril organization”, “bone mineralization” and “mesenchymal cell differentiation” (Figure 3C). As such, c3 may represent progenitor cells closely associated with the fibrosis and pathologic mineralization of adipose tissues in lymphedema. C3-specific regulators such as *KLF13, KLF2* and *JUND* could serve as potential targets for medical intervention (Figure 3D). Cluster c1 was phenotypically close to c3, and its signature was also enriched with extracellular matrix remodeling pathways such as “collagen fibril organization”. Cluster c5 was an ASC subpopulation displaying a unique pattern with a high expression of metallothionein genes such as *MT1X, MT2A, MT1E, MT1G, MT1M* and *MT1A* (Figure 3B). Given that metallothionein proteins mainly play roles in protection against damage associated with heavy metal toxicity, endoplasmic reticulum stress or oxidative stress (Ruttkay-Nedecky et al., 2013; Yang et al., 2015), c5 may represent a stress-responsive subpopulation. Together, we characterized four previously unrecognized subpopulations of ASCs in the SVF of adipose tissue, and found that the lymphedema-associated subpopulation c3 may be related to the fibrosis and pathologic mineralization of adipose tissues in lymphedema.

**Figure 3.**
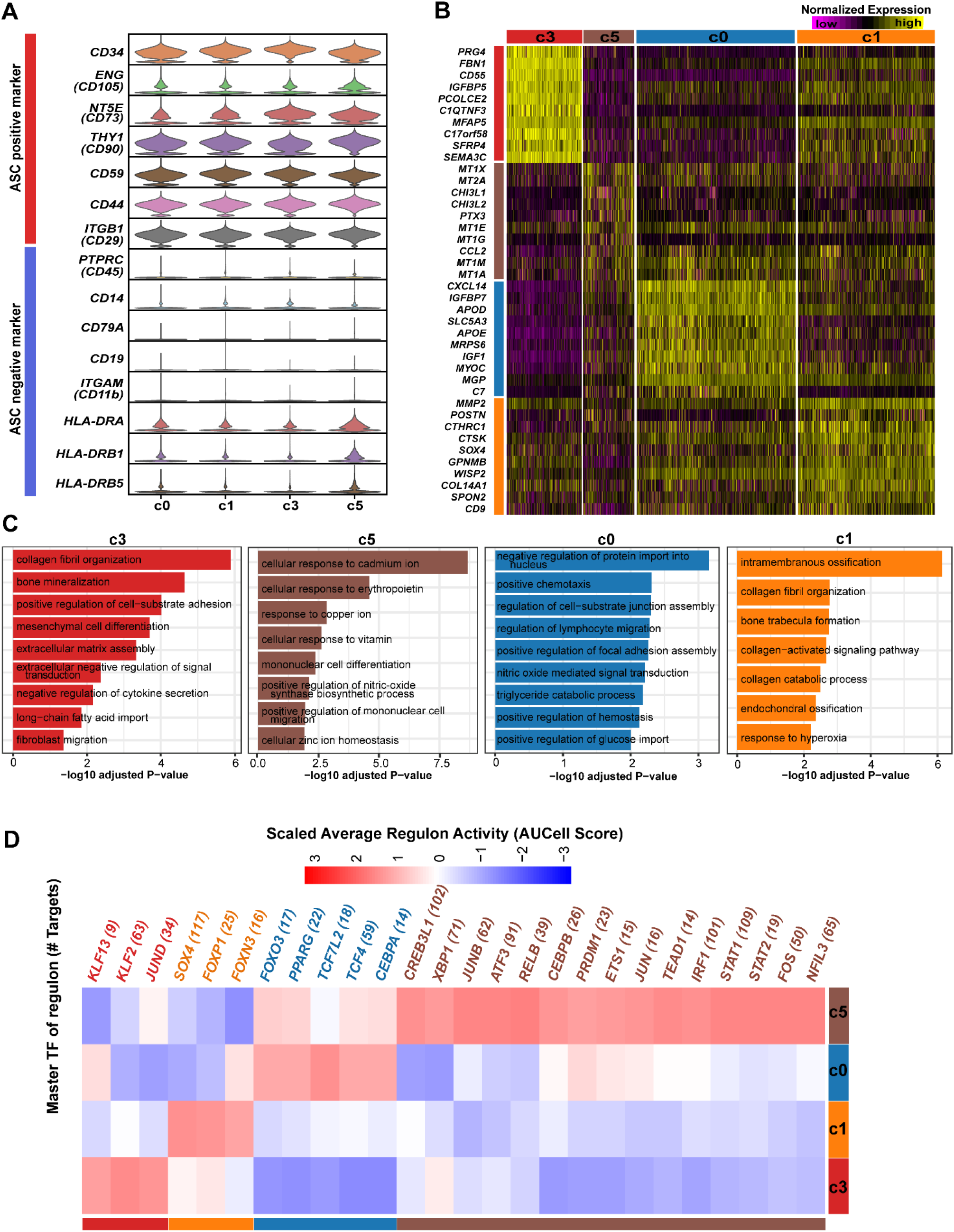
Heterogeneity of ASCs in adipose tissue revealed by single-cell analysis. **(A)** The expression of marker genes normally used for identifying freshly isolated or cultured ASCs. **(B)** Distinct expression profiles displayed by the four subpopulations of ASCs. **(C)** Enriched Gene Ontology terms of the molecular signature for each subpopulation. Adjusted p-value < 0.05. **(D)** Subpopulation-specific regulons of each subpopulation revealed by SCENIC analysis.

### Dysregulated pathways and genes in the ASCs of cancer-related lymphedema

Single-cell RNA-seq allows unbiased analysis of lineage-specific transcriptomic changes in diseased conditions without cell sorting. We next explored the dysregulated pathways through gene set enrichment analysis (GSEA), which facilitates biological interpretation by robustly detecting concordant differences at the gene set or pathway level (Emmert-Streib and Glazko, 2011). Extracellular matrix-related pathways such as “extracellular matrix organization” and “collagen formation” were significantly upregulated (GSEA; FDR q-value < 0.05; Figure 4A; Table S5), which is in line with the fibrosis of adipose tissue in lymphedema. Glycosylation is a common modification of proteins and lipids, which has been implicated in physiological (e.g., cell differentiation) and pathophysiological states (e.g., autoimmunity and chronic inflammation) (Reily et al., 2019). Strikingly, glycosylation-related pathways such as “O-linked glycosylation” and “diseases of glycosylation” were significantly upregulated, which suggests that increased glycosylation or altered glycosylation patterns in ASCs may contribute to pathogenesis. In addition, “SUMOylation of DNA damage response and repair proteins” was upregulated, reflecting DNA damage induced by chronic inflammation (Ioannidou et al., 2016). Compared with the healthy state, ASCs in lymphedema displayed downregulated protein translation, energy metabolism and response to endoplasmic reticulum stress (Figure 4A), reflecting impaired cellular functions at the late stage of lymphedema. Notably, interleukin 10 (IL10) signaling was downregulated in ASCs from lymphedema.

**Figure 4.**
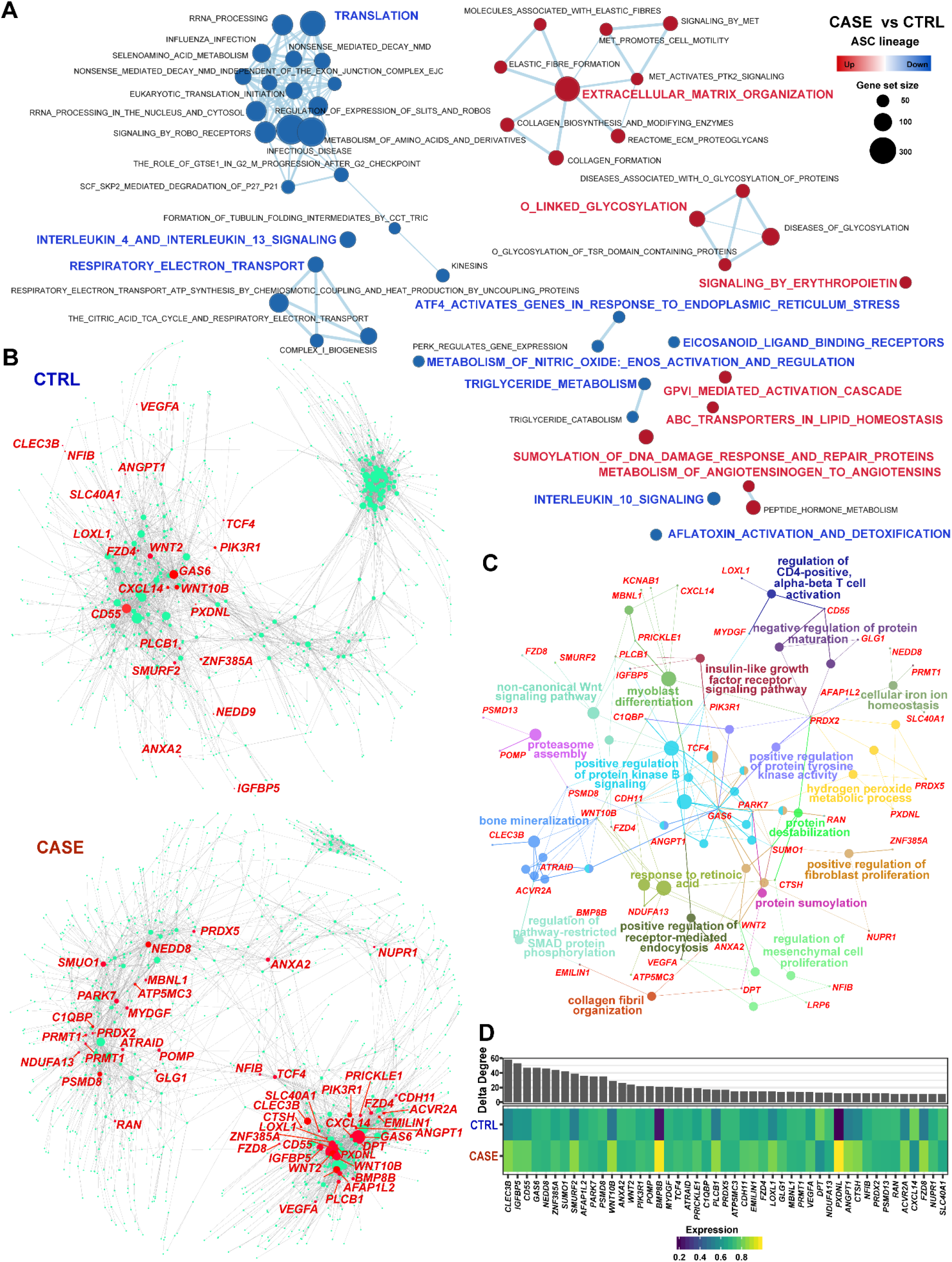
Dysregulated genes and pathways of ASCs in adipose tissue derived from cancer-related lymphedema. **(A)** Gene set enrichment analysis reveals up- and down-regulated pathways of ASCs in CASE versus CTRL. An FDR q-value < 0.05 was considered to be statistically significant. **(B)** Comparative analysis of the gene regulatory networks of ASCs between the CASE (lower panel) and CTRL (upper panel) groups reveals dysregulated genes in ASCs. The node size reflects the degree centrality. The representative genes dysregulated in CASE ranked by delta degree are labeled in red. **(C)** Network view of the functional enrichment for the dysregulated genes shown in B. Small dots denote genes and large nodes represent Gene Ontology terms. The node size represents the number of genes associated with the Gene Ontology term. Adjusted p-value < 0.05. **(D)** Delta degree centrality (upper panel) and average expression across cells in CASE and CTRL (lower panel).

Although the role of IL10 signaling has seldom been discussed in nonimmune cells as targets (Rajbhandari et al., 2018), the downregulation of the expression of *IL10* (Table S5), an important anti-inflammatory cytokine secreted by ASCs, may suggest a reduced immunosuppression capability of ASCs in lymphedema. Unexpectedly, we found decreased adipogenesis for ASCs in lymphedema, as evidenced by the significantly reduced expression of *PPARG* and *CEBPA* (Figure S3A and S3B), the master regulators in adipogenesis (Januszyk et al., 2013), as well as significantly decreased adipogenesis score (Wilcoxon rank sum test p-value < 2.2e-16; Figure S3C). In addition, we found significantly increased osteogenesis of ASCs in lymphedema (Figure S3D), which reflects aberrant differentiation in diseased conditions.

Next, we built gene regulatory networks from single-cell data using a novel method implemented in bigScale2 (Iacono et al., 2019), which allows us to quantify the biological importance of genes and find dysregulated genes in diseased conditions. Figure 4B shows the regulatory networks constructed for ASCs in healthy (upper panel) and diseased conditions (lower panel). Comparative analysis between the two networks revealed a list of genes that were greatly increased in degree centrality (the number of edges connected to a given node; Figure 4B; Table S6) in lymphedema, reflecting their potential roles in the pathogenesis. These genes were mainly involved in bone mineralization, positive regulation of protein kinase B signaling, and regulation of mesenchymal cell proliferation and differentiation (Figure 4C). Notably, *CLEC3B*, encoding a protein implicated in the mineralization process, ranked at the top of the list based on changes in degree centrality (Figure 4D). The expression of *CLEC3B* was upregulated in CASE compared to CTRL (Figure 4D) and was especially high in the lymphedema-associated subpopulation c3 (Table S3), thus highlighting the role of pathologic mineralization of adipose tissues in the pathogenesis of lymphedema. Similarly, the expression of *ZNF385A*, a transcription factor implicated in fibroblast proliferation and differentiation, was also upregulated in CASE (Figure 4D) and was especially high in the lymphedema-associated subpopulation c3.

Together, our results highlight the pathological changes in ASCs, which displayed enhanced fibrosis, mineralization and proliferation as well as compromised immunosuppression capacity, in the severe stage of lymphedema.

### Adipose tissue of lymphedema displays immunological dysfunction characterized by a striking depletion of anti-inflammatory macrophages

Tissue-resident or infiltrated macrophages are phenotypically heterogeneous in a tissue/state-dependent manner (Varol et al., 2015). We next explored the phenotypic differences among the three lymphedema-associated macrophage subpopulations (c6, c8 and c11). These subpopulations displayed distinct expression profiles (Figure 5A; Table S7). Compared with other subpopulations, c6 showed high expression of *LYVE1*, a marker gene associated with tissue-resident macrophages (Lim et al., 2018). It also displayed high expression of markers for M2-polarized (alternatively activated) macrophages, including *RNASE1, SELENOP, MRC1* and *CD163* (Figure 5B), which harbor an antiinflammatory phenotype (Varol et al., 2015). Thus, the *LYVE1^+^* c6 cluster represented a resident-like macrophage subpopulation with an M2 phenotype. Compared with the others, cluster c8 expressed higher levels of *IL1B*, a pro-inflammatory cytokine, and markers for M1-polarized (classically activated) macrophages such as *FCGR1A, TNF* and *FPR2* (Jablonski et al., 2015). The *IL1B* ^high^ cluster c8 thus represented a proinflammatory macrophage subpopulation with an M1 phenotype. Cluster c11 expressed high levels of *CD1C*, encoding an antigen-presenting molecule, and MHC class II genes (e.g., *HLA-DQA1, HLA-DPB1* and *HLA-DPA1;* Figure 5A). It expressed both M1 and M2 markers, e.g., *CD86* and *MRC1*, respectively (Figure 5B). The molecular signature of c11 was enriched with antigen presentation-related terms such as “antigen processing and presentation of exogenous antigen” (Figure 5C). These results suggest that the *CD1C* ^high^ cluster c11 represented a specialized antigen-presenting macrophage subpopulation. Furthermore, we identified subpopulation-specific regulons through SCENIC analysis (Figure 5D), which could serve as potential targets for medical intervention, for example, targeting the key regulators of the proinflammatory macrophage subpopulation c8 (e.g., *CEBPB, FOSL2, STAT1* and *IRF7*).

**Figure 5.**
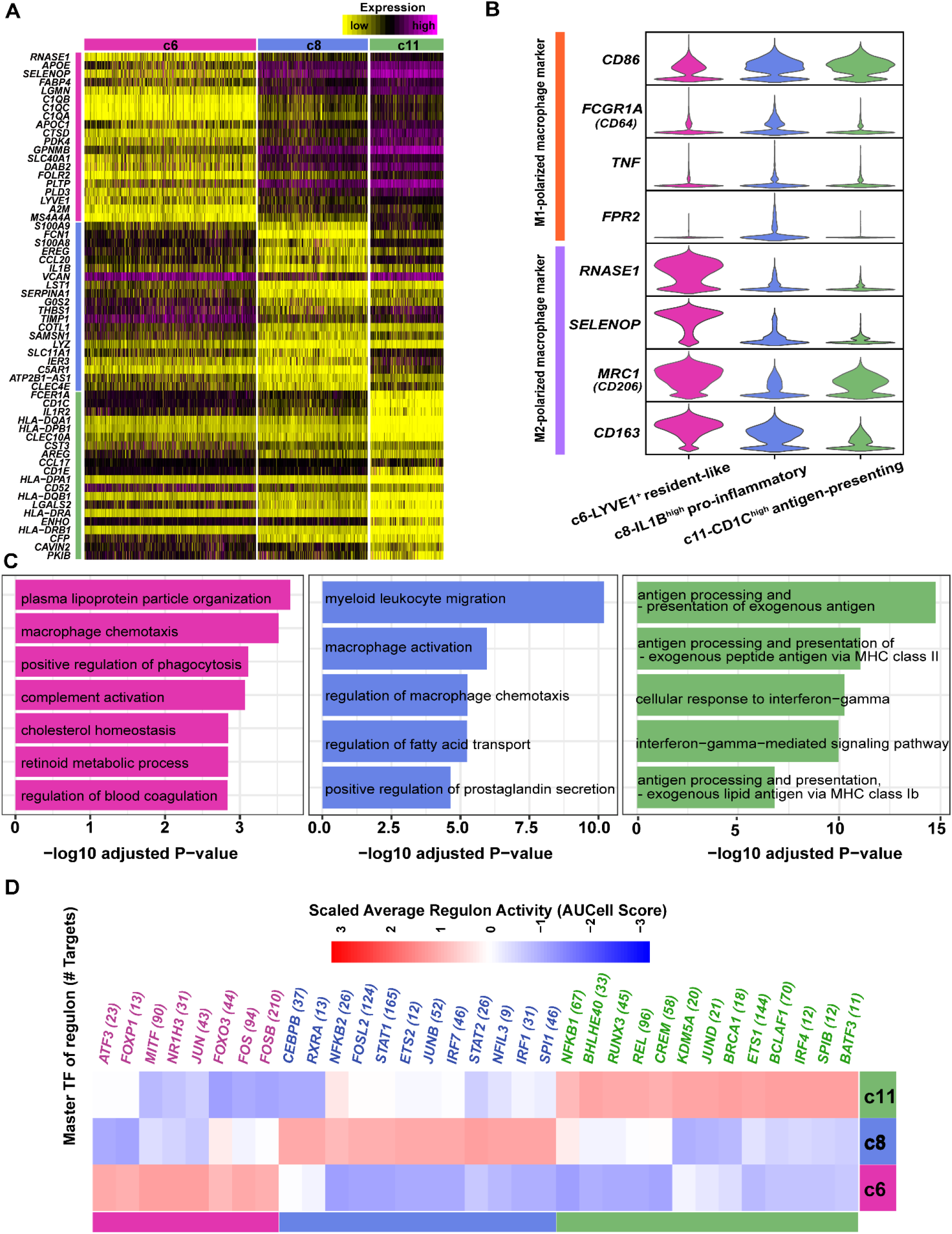
The phenotypic differences among the three lymphedema-associated macrophage subpopulations. **(A)** Distinct expression profiles of the three macrophage subpopulations. **(B)** Expression of M1- or M2-polarized macrophage markers in the three subpopulations. **(C)** Enriched Gene Ontology terms of the molecular signature for each subpopulation. Adjusted p-value < 0.05. **(D)** Subpopulation-specific regulons of each subpopulation revealed by SCENIC analysis.

As mentioned above, the macrophage lineage, especially subpopulation c6, was dramatically reduced in lymphedema (Figure 2B and 2C). We calculated the ratio of c6/c8, as a proxy of the ratio of M1/M2, and found that it was greatly decreased in lymphedema (0.76 in CASE versus 2.03 in CTRL). Together, these results suggest that immunological dysfunction characterized by a striking depletion of anti-inflammatory macrophages occurred in the adipose tissue of lymphedema. Transplantation of *LYVE1^+^* macrophages could thus potentially serve as a cellular therapy for cancer-related lymphedema.

### Cell-cell communication analysis reveals a perivascular ligand-receptor interaction module and communication changes for ASCs in cancer-related lymphedema

The single-cell dataset provided us with a unique chance to analyze cell-cell communication mediated by receptor-ligand interactions. To define the cell-cell communication landscape and uncover its alterations in diseased conditions, we performed analysis using CellPhoneDB 2.0 (Efremova et al., 2019), which contains a curated repository of ligand-receptor interactions and a statistical framework for predicting enriched interactions between two cell types from single-cell transcriptomics data. Strikingly, we identified a densely connected communication network among macrophages, ASCs and vascular endothelial cells in both conditions (Figure 6A), which is concordant with our knowledge that macrophages, especially *LYVE1+* macrophages (Lim et al., 2018), and ASCs (Baer, 2014) are spatially associated with the blood vasculature. In line with this, we found that ASCs were the predominant source of the macrophage colony stimulating factor CSF1 (Figure S4A), which is critical for the survival of tissue macrophages through the activation of the receptor CSF1R (Hume and MacDonald, 2012). The expression of *CSF1* in ASCs was significantly higher in lymphedema than in healthy controls (Figure S4B), reflecting enhanced signals broadcast by ASCs in the diseased state. We therefore identified a perivascular ligand-receptor signal module. Compared with the healthy controls, the total number of interactions for almost all lineages increased in lymphedema (Figure 6A), reflecting enhanced intercellular communications in diseased conditions. Notably, the most abundant interactions in the network occurred between ASCs and macrophages in heathy controls, whereas the most abundant interactions occurred between ASCs and vascular endothelial cells in lymphedema (Figure 6B). Furthermore, we identified the ligand-receptor pairs showing significant changes in specificity between any one of the non-ASC lineages and ASCs in diseased versus healthy conditions (ASCs express receptors and receive ligand signals from other lineages; Figure 6C; Table S9). Notably, PDGFD-PDGFR complex interactions were significantly enhanced between a number of lineages (vascular endothelial cells, mast cells, NKT cells and pericytes) and ASCs in lymphedema. Increased secretion of PDGFD or enhanced PDGFD signaling has been associated with aberrant proliferation and differentiation of mesenchymal cells in a number of diseases such as fibrosis and cancer (Folestad et al., 2018; Wang et al., 2009). Our results suggest that PDGFD signaling may contribute to the enhanced fibrosis and proliferation of ASCs in lymphedema. In addition, we also explored the alterations in ligand signals broadcast by ASCs (Figure 6D). Notably, a number of chemokine signals, including CXCL8, CXCL12 and CCL2, broadcast by ASCs were significantly altered. For example, CXCL12-ACKR3 interactions between ASCs and BCs or DCs become significantly more specific in lymphedema than in healthy conditions (permutation test p-value < 0.05). The CXCL12/CXCR4/ACKR3 axis has been considered a potential therapeutic target for a wide variety of inflammatory diseases, not only by interfering with leukocyte recruitment but also by modulating immune responses (García-Cuesta et al., 2019). Together, the intercellular communication analysis revealed a perivascular signal module in adipose tissue and identified ligand-receptor interaction changes for ASCs in lymphedema, which could serve as potential targets for medical intervention.

**Figure 6.**
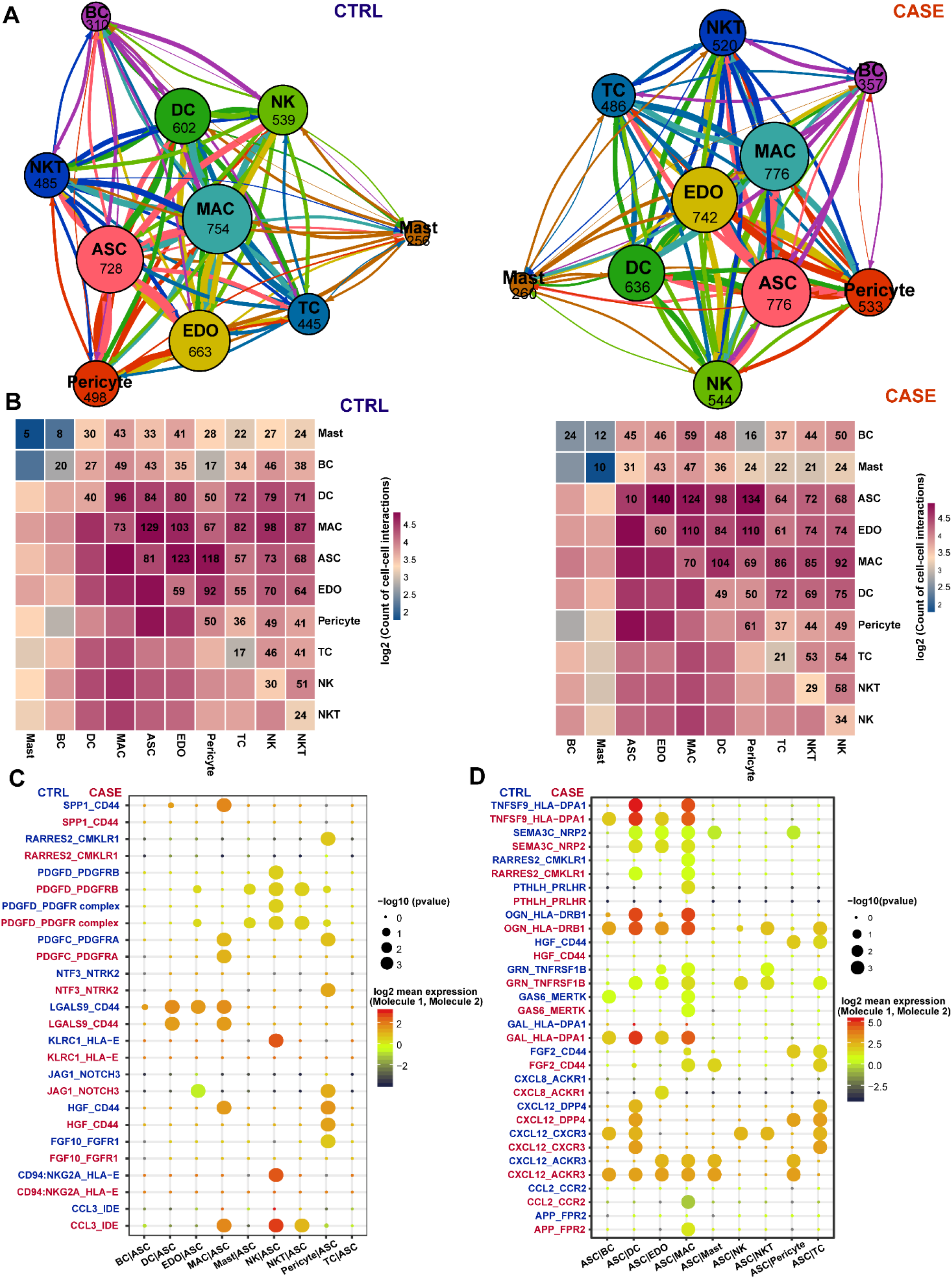
Cell-cell communication analysis reveals a perivascular ligand-receptor interaction module and communication changes for ASCs in cancer-related lymphedema. **(A)** Interlineage communication networks in adipose tissue from patients with lymphedema (CASE; right panel) and healthy people (CTRL; left panel). The total number of communications is shown for each cell lineage. The line color indicates that the ligands are broadcast by the cell lineage in the same color. The line thickness is proportional to the number of broadcast ligands. **(B)** Heatmap shows the number of communications between any two lineages in the CASE (right panel) and CTRL (left panel) groups. **(C)** The ligand-receptor pairs that were shown significant changes in specificity between any one of the non-ASC lineages and ASCs in CASE versus CTRL. ASCs express receptors and receive ligand signals from other lineages. The dot size reflects the p-value of the permutation tests for lineage-specificity. The dot color denotes the mean of the average ligand-receptor expression in the interacting lineages. **(D)** The ligand-receptor pairs that were shown significant changes in specificity between ASCs and any one of the non-ASC lineages in CASE versus CTRL. ASCs express ligands and broadcast ligand signals for other lineages. ASC: adipose-derived stromal/stem/progenitor cell; BC: B cell; DC: dendritic cell; EDO: endothelial cell; MAC: macrophage; NK: natural killer cell; NKT: natural killer T cell; TC: T cell.

### Mapping the previously reported candidate genes predisposing to cancer-related lymphedema to cell subpopulations in the SVF

Genetic susceptibility may partially explain the development of secondary lymphedema in cancer survivors (Newman et al., 2012). The single-cell RNA-seq dataset provided us an unpreceded chance to map the previously reported 18 candidate genes predisposing to cancer-related lymphedema (Visser et al., 2019) to cell subpopulations in the SVF. As shown in Figure 7, most predisposing genes were highly expressed in a specific cell subpopulation, including *HGF, MET, GJC2, IL1A, IL4, IL6, IL10, IL13, NRP2, VCAM1, FOXC2, KDR, FLT4* and *RORC.* Notably, *GJC2*, the most likely causal gene (Visser et al., 2019), was highly expressed in the lymphedema-associated ASC subpopulation c3. The expression of four candidate genes, including *MET, KDR, FLT4* and *FOXC2*, was highly specific in vascular endothelial cells (c13) or pericytes (c20), reflecting the role of vascular cells in the pathogenesis. Together, our results will help elucidate the cellular and molecular mechanisms underlying the pathogenesis of cancer-related lymphedema.

**Figure 7.**
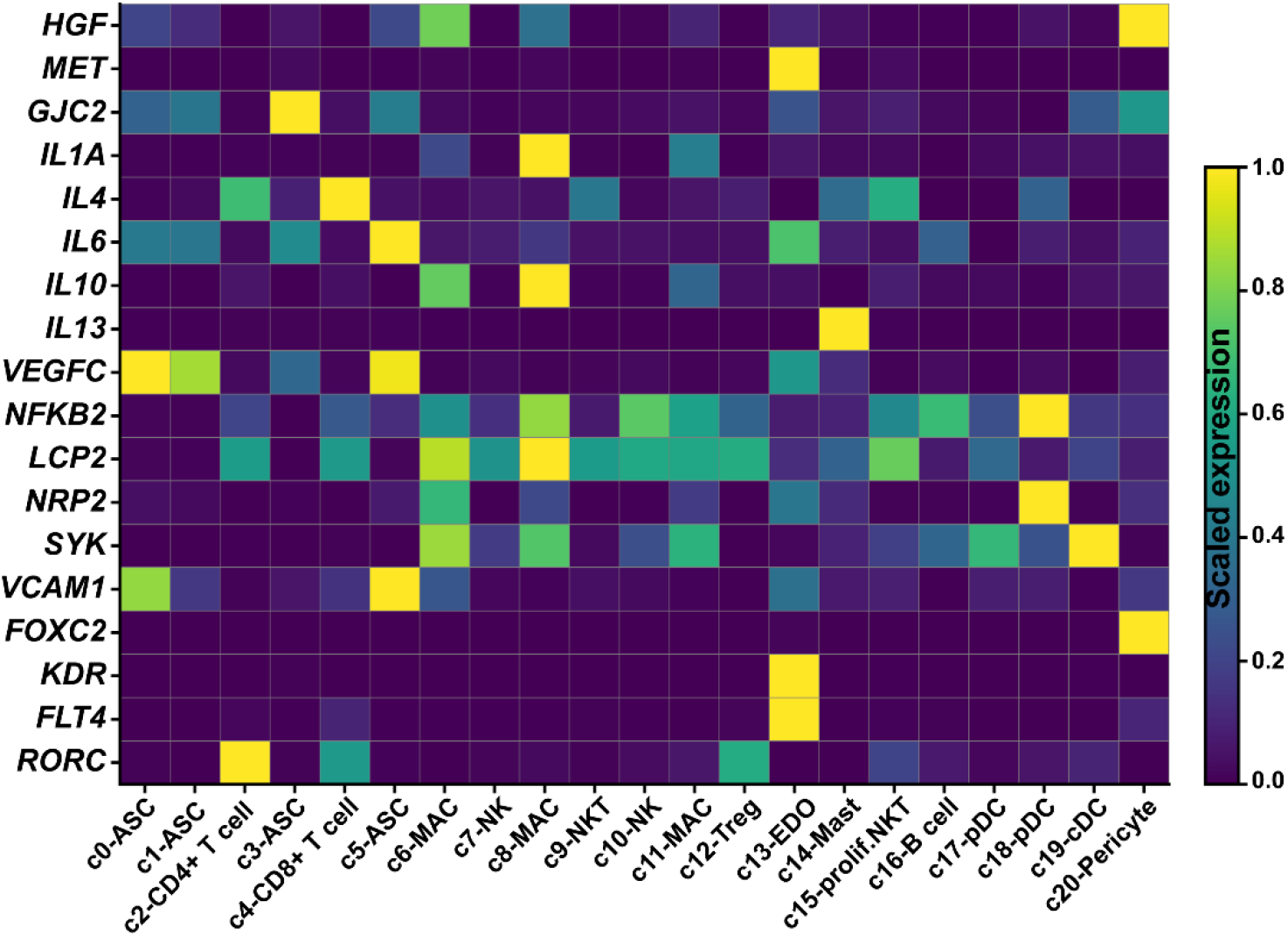
The expression of 18 previously reported candidate genes predisposing to cancer-related lymphedema in each cell cluster.

## Discussion

Understanding the cellular heterogeneity and regulatory changes of tissues in diseased conditions is fundamental to successful medical therapy development. Here, we performed single-cell RNA-seq of 70,209 cells of the SVF of subcutaneous adipose tissue from patients with cancer-related lymphedema and healthy donors. Unbiased clustering revealed 21 cell clusters, which were assigned to 10 cell lineages. One of the four ASC subpopulations, c3, was significantly expanded in lymphedema. Functional analysis revealed that this lymphedema-associated ASC subpopulation may be related to the fibrosis and pathologic mineralization of adipose tissues in lymphedema. We also identified c3-specific regulators, such as *KLF13, KLF2* and *JUND*, which could serve as potential targets for medical intervention. Dysregulated pathways and genes of ASCs in lymphedema were identified through GSEA and differential regulatory network analysis, which reflect the pathophysiological changes in ASCs in lymphedema: enhanced fibrosis, mineralization and proliferation as well as compromised immunosuppression capacity. In addition, we characterized the three subpopulations of macrophages, and found that the adipose tissue of lymphedema displayed immunological dysfunction characterized by a striking depletion of antiinflammatory macrophages, i.e., *LYVE^+^* resident-like macrophages. Cell-cell communication analysis revealed a perivascular ligand-receptor interaction module among ASCs, macrophages and vascular endothelial cells in adipose tissue. Finally, we mapped the previously reported candidate genes predisposing to cancer-related lymphedema to cell subpopulations in SVF.

Lymphedema is characterized by excess adipose deposition in the affected limbs (Mehrara and Greene, 2014); however, the underlying mechanism remains elusive. Previous studies suggested enhanced adipogenesis, i.e., the differentiation of adipocytes from ASCs in mouse models (Aschen et al., 2012) and human patients (Januszyk et al., 2013), based on a limited number of marker genes. In contrast, our large-scale single-cell analysis did not find any significantly upregulated pathways associated with adipogenesis. Instead, we found that ASCs from lymphedema may have decreased adipogenesis (Figure S3) and enhanced proliferation ability (Figure S2). The enhanced proliferation of ASCs from lymphedema is consistent with the findings of a study based on bulk RNA-seq (Xiang et al., 2020). Histological evidence has shown that hypertrophic (cell enlargement) adipocytes are frequently observed, especially in the severe stages of lymphoedema (Tashiro et al., 2017). Therefore, we think that the excess adipose deposition may be mostly attributed to the enhanced proliferation ability of ASCs and cell enlargement of adipocytes at least in the severe stage of lymphoedema.

Stage III lymphedema, also known as lymphostatic elephantiasis, is a severe condition in which the tissue becomes extremely swollen, thickened and fibrotic (hardened) (Lawenda et al., 2009). Concordant with the enhanced fibrosis, we found that extracellular matrix-related pathways, such as “extracellular matrix organization” and “collagen formation”, were significantly upregulated in ASCs from lymphedema (Figure 4A). In addition, differential regulatory network analysis revealed that the genes involved in the bone mineralization process, e.g., *CLEC3B*, ranked at the top based on the changes in degree centrality (Figure 4D). We also found significantly increased osteogenesis scores based on a set of osteogenesis-related genes in ASCs from lymphedema (Figure S3D). Furthermore, we pinpointed the ASC subpopulation closely associated with lymphedema, i.e., c3, which was significantly expanded in lymphedema. The molecular signature of this subpopulation was enriched with pathways such as “collagen fibril organization” and “bone mineralization” (Figure 3C), suggesting that this subpopulation was related to both the fibrosis and pathologic mineralization of adipose tissues in lymphedema. Altogether, our results indicated that the hardened tissue at the severe stage of lymphoedema may not only be attributed to fibrosis, but also to pathologic mineralization of adipose tissues, which has not been recognized before. Pathological mineralization occurs in nearly all soft tissues and is associated with diverse human diseases such as cancer and atherosclerosis, but is sometimes overlooked (Tsolaki and Bertazzo, 2019). Our study highlights the aberrant differentiation or pathological mineralization of ASCs in lymphoedema, which may serve as a novel angle for treatment.

We found a striking depletion of antiinflammatory macrophages, i.e., the c6 *LYVE1^+^* resident-like subpopulation, in the adipose tissue of lymphedema (Figure 2C; Figure4B). It has been reported that *LYVE1^+^* macrophages contribute to the homeostasis of the aorta through the control of collagen deposition by smooth muscle cells, thus preventing arterial stiffness (Lim et al., 2018). In addition, our analysis revealed a perivascular ligand-receptor interaction module among ASCs, macrophages and vascular endothelial cells in adipose tissue (Figure 6), and found that ASCs were the predominant source of the macrophage colony stimulating factor CSF1 (Figure S4A). These results reflect the close relationship between macrophages and ASCs in adipose tissue. The depletion of macrophages may contribute to the pathological changes in ASCs in lymphedema. Previous studies have proven that targeting immune cell subpopulations, such as CD4^+^ helper T cells (Zampell et al., 2012a), was effective for alleviating the effects of lymphedema. We therefore propose that transplantation of *LYVE^+^* resident-like anti-inflammatory macrophages could serve as a cellular therapy for cancer-related lymphedema. Since the expression of *CSF1* in ASCs was even significantly higher in lymphedema than in healthy controls (Figure S4B), we reason that the mechanism underlying the depletion of macrophages, especially for the *LYVE1^+^* macrophages, may not be due to pathological changes in ASCs. However, the precise mechanism remains to be explored.

In conclusion, we provided the first comprehensive analysis of cellular heterogeneity, lineage-specific regulatory changes, and intercellular communications of the SVF in adipose tissues from cancer-related lymphedema at a single-cell resolution. Our study revealed lymphedema-associated cell subpopulations and dysregulated pathways in ASCs, as well as a strong depletion of *LYVE^+^* anti-inflammatory macrophage in lymphedema, which could serve as potential targets for medical therapies. Our large-scale dataset constitutes a valuable resource for further investigations of the mechanism of cancer-related lymphedema.

## Methods

### Ethics approval

All human patient recruitments and tissue sampling procedures complied with the ethics regulations approved by Peking Union Medical College Hospital. Each subject provided written informed consent.

### Specimen preparation and SVF Isolation

Adipose tissue specimens were obtained from the affected thighs of five female patients with secondary lymphoedema (stage III) following surgical intervention for cervical cancer. As a control group, liposuction specimens from the thighs of four healthy female donors were collected during surgery for cosmetic purposes. All fresh specimens were subjected to SVF isolation. Briefly, each specimen was washed several times with Hank’s balanced salt solution (HBSS). Then, it was digested with 0.15% collagenase supplied with 4% penicillin streptomycin solution (P/S) at 37°C for 30 minutes. Subsequently, high-glucose Dulbecco’s Modified Eagle’s Medium (DMEM) with 10% fetal bovine serum (FBS) was added, and the sample was centrifuged at 4°C for 10 minutes. The pellet was resuspended in high-glucose DMEM with 10% FBS, filtered through a 100-μm strainer, and then centrifuged at 4 °C for 5 minutes. The obtained cell suspensions were resuspended in HBSS, and red blood cell lysis buffer was added. Then, it was centrifuged again, resuspended in HBSS with 0.04% bovine serum albumin (BSA) and filtered through a 40-μm strainer. Finally, the cells were centrifuged and resuspended in Dulbecco’s Phosphate Buffered Saline (DPBS).

### Single-cell RNA-seq library preparation and sequencing

Single-cell Gel Beads-in-Emulsion (GEM) generation, barcoding, post GEM-RT cleanup, cDNA amplification and cDNA library construction were performed using Chromium Single Cell 3’ Reagent Kit v3 chemistry (10X Genomics, USA) following the manufacturer’s protocol. The resulting libraries were sequenced on a NovaSeq 6000 system (Illumina, USA).

### Sample demultiplexing, barcode processing and UMI counting

The official software Cell Ranger v3.0.2 (https://support.10xgenomics.com) was applied for sample demultiplexing, barcode processing and unique molecular identifier (UMI) counting. Briefly, the raw base call files generated by the sequencers were demultiplexed into reads in FASTQ format using the “cellranger mkfastq” pipeline. Then, the reads were processed using the “cellranger count” pipeline to generate a gene-barcode matrix for each library. During this step, the reads were aligned to the mouse human reference genome (version: GRCh38). The resulting gene-cell UMI count matrices of all samples were ultimately concatenated into one matrix using the “cellranger aggr” pipeline.

### Data cleaning, normalization, feature selection, integration and scaling

The concatenated gene-cell barcode matrix was imported into Seurat v3.1.0 for data preprocessing. To exclude genes likely detected from random noise, we filtered out genes with counts in fewer than 3 cells. To exclude poor-quality cells that might have resulted from doublets or other technical noise, we filtered cell outliers (> third quartile + 1.5 × interquartile range or < first quartile – 1.5 × interquartile range) based on the number of expressed genes, the sum of UMI counts and the proportion of mitochondrial genes. To further remove doublets, we filtered out cells based on the predictions by Scrublet (Wolock et al., 2019). In addition, cells enriched in hemoglobin gene expression were considered red blood cells and were excluded from further analyses. The sum of the UMI counts for each cell was normalized to 10,000 and log-transformed. For each sample, 2,000 features (genes) were selected using the “FindVariableFeatures” function of Seurat under the default settings. To correct for potential batch effects and identify shared cell states across datasets, we integrated all the datasets via canonical correlation analysis (CCA) implemented in Seurat. To mitigate the effects of uninteresting sources of variation (e.g., cell cycle), we regressed out the mitochondrial gene proportion, UMI count, S phase score and G2M phase score (calculated by the “CellCycleScoring” function) with linear models using the “ScaleData” function. Finally, the data were centered for each gene by subtracting the average expression of that gene across all cells, and were scaled by dividing the centered expression by the standard deviation.

### Dimensional reduction and clustering

The expression of the selected genes was subjected to linear dimensional reduction through principal component analysis (PCA). The first 30 components of the PCA were used to compute a neighborhood graph of the cells. The neighborhood graph was ultimately embedded in two-dimensional space using the nonlinear dimensional reduction method of uniform manifold approximation and projection (UMAP) (Becht et al., 2019). The neighborhood graph of cells was clustered using Louvain clustering (resolution=0.6) (Blondel et al., 2008).

### Differential expression and functional enrichment analysis

Differentially expressed genes between two groups of cells were detected with the likelihood-ratio test (test.use: “bimod”) implemented in the “FindMarkers” function of Seurat. The significance threshold was set to an adjusted p-value < 0.05 and a log2-fold change > 0.25. Functional enrichment analyses of a list of genes were performed using ClueGO (Bindea et al., 2009) with an adjusted p-value threshold of 0.05.

### Gene set enrichment analysis

All the expressed genes were preranked by Signal2Noise (the difference in means between CASE and CTRL scaled by the standard deviation). Then, the ranked gene list was imported into the software GSEA (version: 4.0.1) (Subramanian et al., 2005). An FDR q-value < 0.05 was considered to be statistically significant. Precompiled gene sets, i.e., REACTOME pathways in MSigDB (version: 7.0) (Liberzon et al., 2015) were used in this analysis. The results were visualized using the EnrichmentMap plugin of Cytoscape (version: 3.7.0).

### Differential proportion analysis

To determine whether the change in the cell proportion of a specific lineage or cluster compared with the control was expected by chance, we performed a permutation-based statistical test (differential proportion analysis; DPA) as described previously (Farbehi et al., 2019). A Bonferroni-corrected p-value < 0.05 was considered to be statistically significant.

### Differential regulatory network analysis based on single-cell transcriptomes

Gene regulatory networks were constructed from single-cell datasets and compared using the method implemented in bigScale2 (Iacono et al., 2019). Briefly, gene regulatory networks for the CASE and CTRL were inferred with the ‘compute.network’ function (‘clustering-direct’, quantile.p = 0.90) separately. Genes encoding ribosomal proteins or mitochondrial proteins were excluded from this analysis. Then, the number of edges was homogenized throughout the obtained networks using the ‘homogenize.networks’ function. Finally, changes in node centralities (the relative importance of genes in the network) in the CASE compared to the CTRL group were identified using the ‘compare.centrality’ function. Four measures of centrality, namely degree, betweenness, closeness and pagerank, were considered. The networks were ultimately visualized with Cytoscape (version: 3.7.0).

### Subpopulation-specific regulon analysis

To identify the master regulators driving the cellular heterogeneity among subpopulations, we performed regulon analysis using the R package SCENIC (Aibar et al., 2017). Briefly, coexpression modules were identified, which included a set of genes coexpressed with regulators. Then, only the modules with significant motif enrichment of the regulators were retained, which were referred to as regulons. The activity of each regulon was ultimately scored for each cell. Subpopulation-specific regulons could be found based on the average regulon activity scores of cells in the subpopulation.

### Cell-cell communication analysis based on single-cell transcriptomes

To analyze cell-cell communication based on single-cell transcriptomic datasets, we used CellPhoneDB 2.0 (Efremova et al., 2019), which contains a curated repository of ligand-receptor interactions and a statistical framework for inferring lineage-specific interactions. Briefly, potential ligand-receptor interactions were established based on the expression of a receptor by one lineage and a ligand by another. Only ligands and receptors expressed in greater than 10% of the cells in any given lineage were considered. The labels of all cells were randomly permuted 1000 times and the means of the average ligand-receptor expression in the interacting lineages were calculated, thus generating a null distribution for each ligand-receptor pair in each pairwise comparison between lineages. Ultimately, a p-value for the likelihood of lineage specificity for a given ligand-receptor pair was obtained.

## Author contributions

X. Liu analyzed the data, interpreted the results and wrote the manuscript. M. Y. and Q. X. performed tissue dissociation and library preparation, and participated in drafting the manuscript. Z. L., J. C., J. H. and N. Y. prepared the samples and contributed to the result interpretation. X. Long and Z. Z. conceived the project. W. C. participated in the design of the project.

## Acknowledgments

This work was supported by the grants from the Natural Science Foundation of China (81870229, 81900282) and the Chinese Academy of Medical Sciences Initiative for Innovative Medicine Grant (2016-I2M-1-016).

## Supplemental Materials

**Figure S1. Expression of markers for CD4^+^ T cells, Treg cells, CD8^+^ T cells, proliferation and cytotoxicity in the clusters of T cells and NKT cells.**

**Figure S2. The distribution of cycling scores of ASCs in CASE and CTRL. The cycling score is defined as the sum of the expression of a group of cycling genes.**

**Figure S3. Decreased adipogenesis and increased osteogenesis of ASCs in lymphedema. (A)** The expression of *PPARG*, the master regulator of adipogenesis, was significantly decreased in CASE compared to CTRL. **(B)** The expression of *CEBPA*, another key regulator of adipogenesis, was significantly decreased in CASE compared to CTRL. **(C)** Decreased adipogenesis in CASE compared to CTRL. The adipogenesis score is defined as the sum of the expression of a curated list of genes involved in adipogenesis, including *ACACA, ANGPTL4, APOE, CD36, CEBPA, CEBPB, CEBPD, FASN, INSR, PPARG, SREBF1, IGF1, PLIN2, ADIPOQ, AOC3, AQP7, CITED1, FABP4, LEP, LPL, PCK1, SCD, SLC27A1, SLC2A4, SLCO2A1* and *UCP1*. **(D)** Increased osteogenesis in CASE compared to CTRL. The osteogenesis score is defined as the sum of the expression of a curated list of genes involved in osteogenesis, including *BMP2, COL11A1, COL9A2, COMP, FGFR3, HAPLN1, IHH, PTCH1, SOX5, SOX6, SOX9, TNFSF11, WNT11, WNT4, ACAN, BMP7, CD151, COL10A1, COL2A1, COL4A1, COL9A3, DMP1, EPYC, IBSP, MEF2C, MMP3, PAPLN, PRG4, RUNX3*, and *MIA.*

**Figure S4. ASCs are the predominant source of the macrophage colony stimulating factor CSF1. (A)** ASCs predominately express *CSF1* (left panel) and macrophages express the receptor *CSF1R* (right panel). **(B)** The expression of *CSF1* in ASCs (left panel) and *CSF1R* in macrophages (right panel) in diseased and healthy states.

**Table S1. Clinical information of the subjects and sequencing quality metrics of the samples.**

**Table S2. Molecular signature for each of the 21 cellular clusters.**

**Table S3. Molecular signature for each subpopulation of ASCs.** The molecular signature was obtained by differential expression analysis between one subpopulation and the others.

**Table S4. ASC subpopulation-specific regulons and their targets revealed by SCENIC analysis.**

**Table S5. Dysregulated pathways of ASCs in cancer-related lymphedema revealed by gene set enrichment analysis.**

**Table S6. Results of node centrality comparisons between the gene regulatory networks of the ASCs in CASE and CTRL.**

**Table S7. Molecular signature for each subpopulation of macrophages. The molecular signature was obtained by differential expression analysis between one subpopulation and the others.**

**Table S8. Macrophage subpopulation-specific regulons and their targets revealed by SCENIC analysis.**

**Table S9. Statistical inference of receptor-ligand specificity between all cell lineages with CellPhoneDB.**

