## Supplementary figures and images for "Single-cell RNA-seq of the stromal vascular fraction of adipose tissue reveals lineage-specific changes in cancer-related lymphedema"

### Figure S1

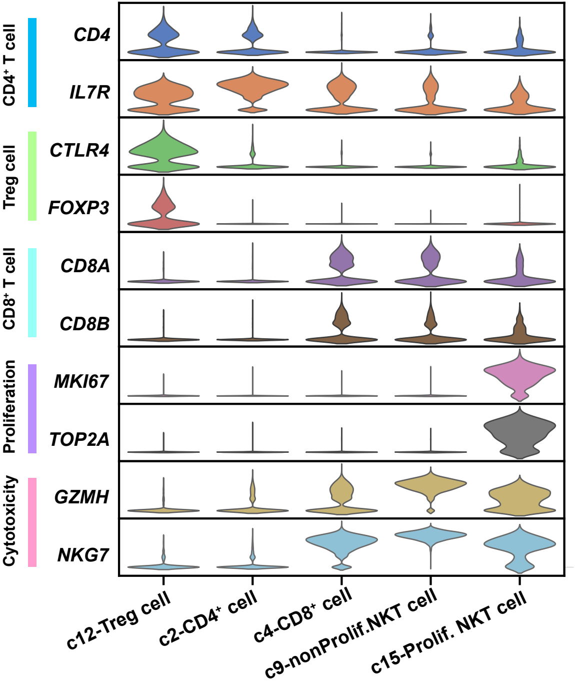

### Figure S2

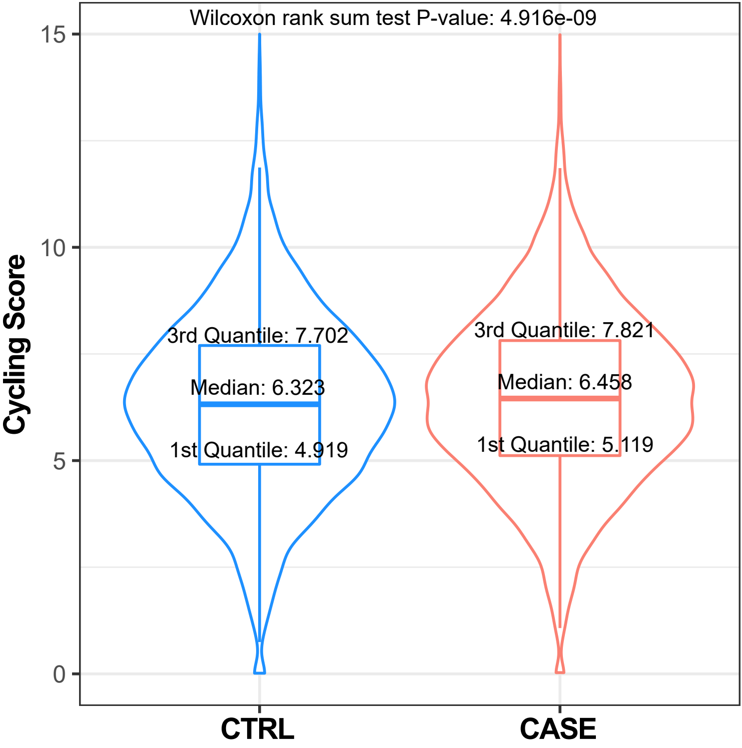

### Figure S3

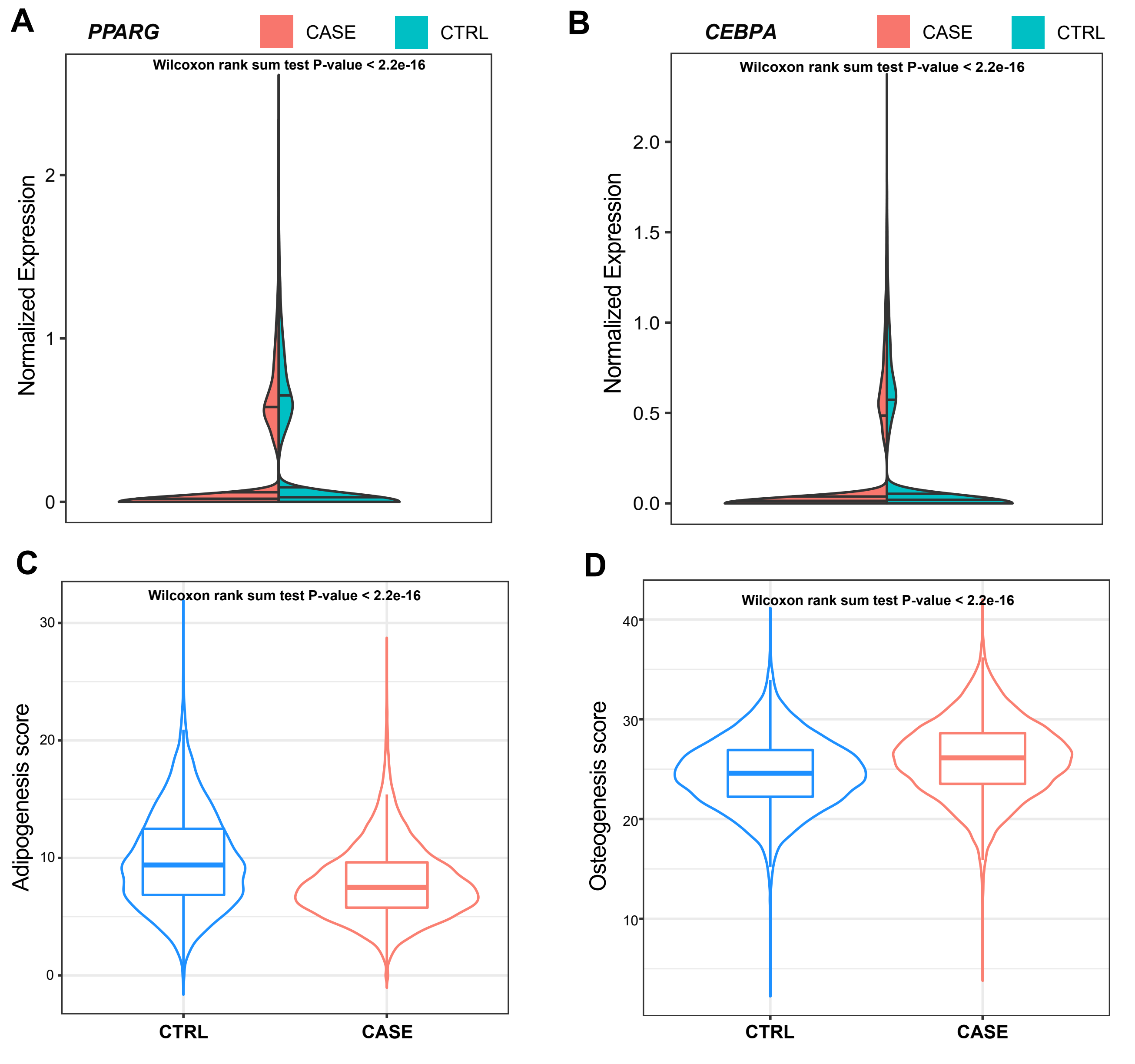

### Figure S4

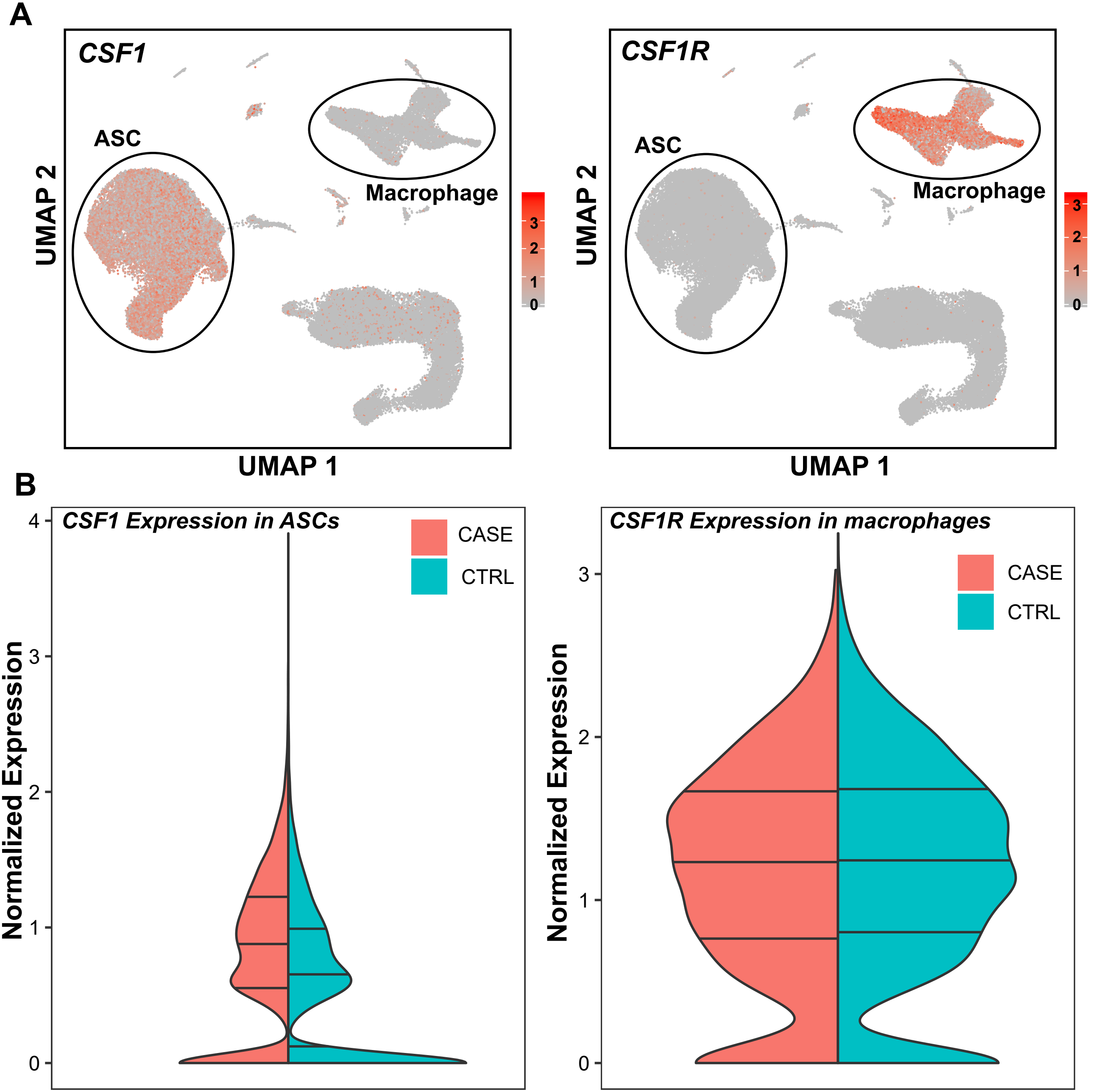
